# Overcoming off-target optical stimulation-evoked cortical activity in the mouse brain *in vivo*

**DOI:** 10.1101/2024.08.14.607152

**Authors:** Simon Weiler, Mateo Vélez-Fort, Troy W. Margrie

**Affiliations:** Sainsbury Wellcome Centre for Neuronal Circuits and Behavior, University College London, 25 Howland Street, London W1T 4JG, United Kingdom

**Keywords:** Optogenetic, Neuropixels, Retinal activation, Laser iIllumination, Cortical activation

## Abstract

Genetic engineering of exogenous opsins sensitive to a wide range of light wavelengths allows the interrogation of brain circuits to an unprecedented temporal and spatial precision. In particular, red-shifted opsins offer access deeper within the brain tissue. It is however crucial to consider the potential unintended back-activation of endogenous opsins due to laser light striking the back of the retina. Here, we found that in complete darkness and with no expression of exogenous opsins, optical fiber laser stimulation at wavelengths of 637 nm (red), 594 nm (orange), or 473 nm (blue) from within the ipsilateral mouse visual cortex resulted in a strong neuronal response in its contralateral counterpart. This neuronal activation occurred even at low laser intensities (1mW at the fiber tip, 31.8mW/mm2) and was most pronounced using red wavelengths. We therefore took advantage of retinal light adaptation using external illumination with a relatively dim ambient light source (20 lux) which was found to completely abolish orange and blue laser-evoked neuronal activation from within the brain, even at high laser intensities (15mW, 477.3mW/mm^2^). To prevent red laser-evoked retinal activation, however, only much lower intensities (2.5mW, 79.6mW/mm^2^) combined with external illumination (20 lux) could be used. These findings demonstrate the critical need for careful selection of light wavelengths and intensities for laser stimulation during optogenetic experiments in the mouse brain *in vivo*. Additionally, light adaptation of the retina through ambient light exposure offers an effective solution to minimize unintended retinal activation.

## 1 Introduction

Optogenetic tools have revolutionized the field of *in vivo* systems neuroscience by offering temporally precise, reversible and cell type-specific modulation of neural activity in the brain (Nagel *et al*. 2003; Boyden *et al*. 2005; Scanziani and Häusser 2009; Fenno, Yizhar, and Deisseroth 2011; Packer, Roska, and Häusser 2013; Emiliani *et al*. 2022). For example the selective activation or silencing of populations of cells expressing genetically encoded wavelength-specific opsins has revealed important causal roles of neuronal circuits underlying animal behaviors (Huber *et al*. 2008; Tsai *et al*. 2009). More recently, short-(473 nm) and long-wavelength (594 nm and 637 nm) opsins have been used to independently modulate the activity of non-overlapping neuronal populations within the same brain area (Klapoetke *et al*. 2014; Hooks 2018; Bauer *et al*. 2021) and expression of dual-color opsins even enable both activation and suppression of activity within the same neurons (Vierock *et al*. 2021). In combination with the use of optical fiber implants these tools allow for spatially restricted stimulation of neurons in deep brain areas (Tsakas *et al*. 2021; Emiliani *et al*. 2022; Adelsberger *et al*. 2014) and thus the dissection of the role of excitation and inhibition within specific circuits during behavior.

Although the delivery of light from within the brain allows the interrogation of circuits in deep structures, it is crucial to consider the non-specific effects of light propagation through brain tissue when performing optogenetic experiments - in particular, when using high light intensities and red-shifted wavelengths. For example, electroretinogram (ERG) recordings indicate that the retina can be directly activated by red, orange and blue light emitted from the tip of the optical fiber implanted within the brain at laser light intensities found to disrupt behavioral tasks in mice (Danskin *et al*. 2015). From a behavioral standpoint such off-target retinal activation-induced artifacts may be overcome using light adaptation that decreases the sensitivity of the retina to wavelengths of light used for optogenetic stimulation (Danskin *et al*. 2015). Importantly, and despite rodents being considered dichromats (highest sensitivity at ultraviolet and green light), off-target activation of the retina has been found to be most prominent when using red light (Danskin *et al*. 2015). This is explained by recent evidence showing that rodents are not red-light blind (Niklaus *et al*. 2020), but also by the increased penetration of long-wavelength light through brain tissue when compared to short-wavelengths (Yaroslavsky *et al*. 2002; Yizhar *et al*. 2011; Jacques 2013; Danskin *et al*. 2015; Lehtinen, Nokia, and Takala 2022).

While the risk of off-target activation of retinal opsins has been shown to lead to behavioral artifacts (Danskin *et al*. 2015), its impact on distant downstream neuronal circuits remains unknown. Moreover, the extent to which retinal light-adaptation using external ambient illumination impacts downstream brain activity is largely unexplored. In this study, we tested the wavelength- and intensity-dependent effect of retinal activation by optical stimulation within the brain on cortical neuronal responses using Neuropixels recordings in the absence of expression of any exogenous opsins. We find that while red, orange and blue light emitted deep within the cortex can cause pronounced modulation of contralateral visual cortical activity this may be ameliorated by the presence of an external ambient light source.

## 2 Results

### In complete darkness and in the absence of exogenous opsins optical-fiber illumination within the brain activates the mouse visual cortex

We first tested whether laser stimulation of the visual cortex at power intensities/irradiances (1 / 31.8; 2.5 / 79.6; 5 / 159.1; 10 / 318.2; 15 / 477.3mW /mW/mm2) and wavelengths (637, 594 and 473 nm) regularly used for *in vivo* optogenetic experiments (Aravanis *et al*. 2007; Fenno, Yizhar, and Deisseroth 2011; Madisen *et al*. 2012; Olsen *et al*. 2012; Pinto *et al*. 2013; Chuong *et al*. 2014; Danskin *et al*. 2015) had an effect on neuronal activity in the absence of genetically-expressed opsins (Danskin *et al*. 2015). For this, we implanted a small-diameter optic fiber in deep layers of the visual cortex (n = 4 mice) and performed Neuropixels recordings in the contralateral visual cortex of head-fixed awake mice under conditions of total darkness (distance between Neuropixels and optic fiber: 4mm, Fig. 1A). Importantly, to avoid direct external stimulation of the retina via light emission from the fiber, the ferrule sleeve of the optical fiber was shielded with blackout material. To prevent light being emitted through the brain and skull, black dental cement was used to seal the craniotomy within and around the 1 × 1.2 cm headplate (Araragi, Alenina, and Bader 2022). Finally, two eye cups were also positioned over the two eyes (Fig. 1A).

**Figure 1.**
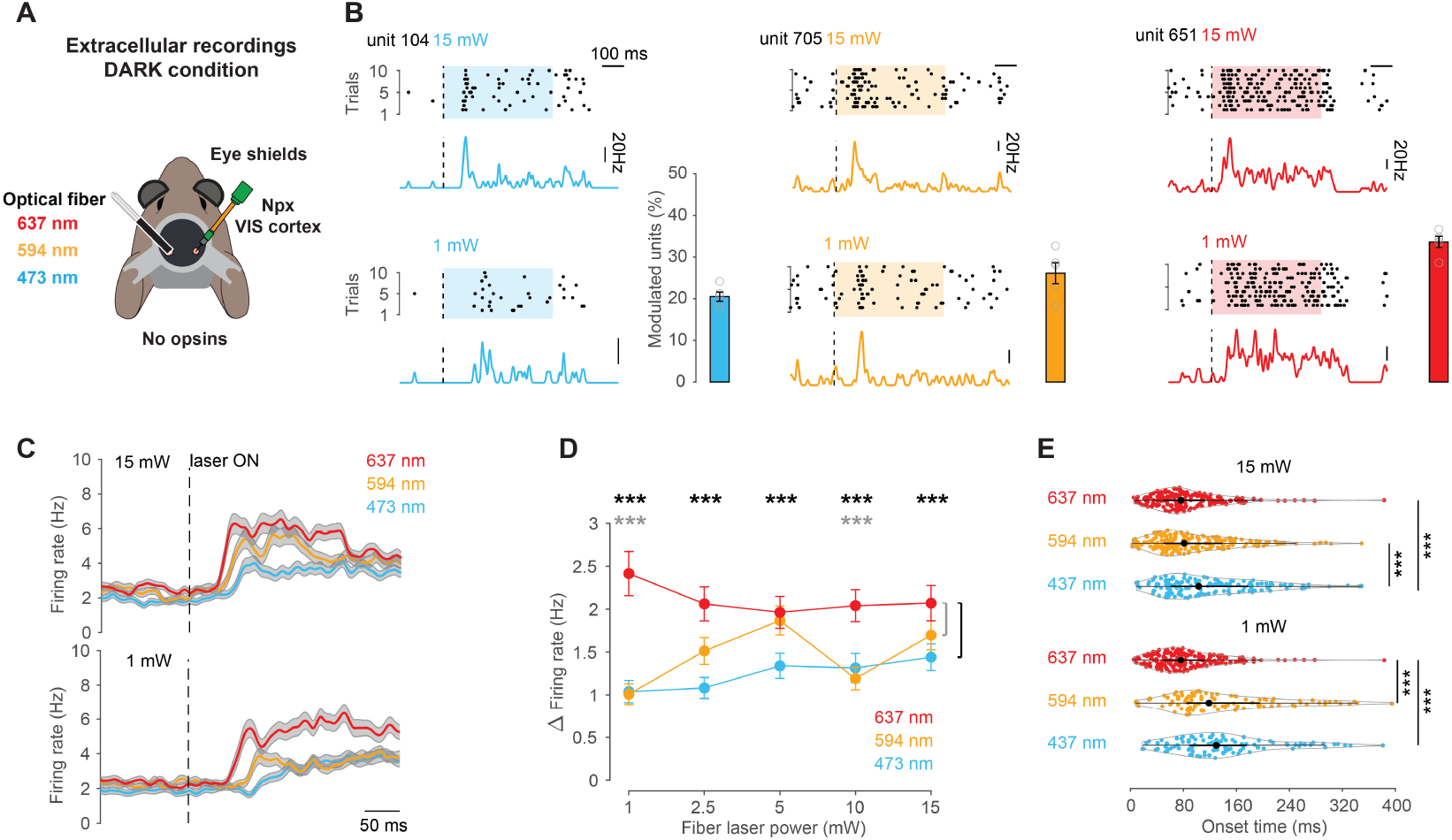
Optical-fiber illumination in the brain using different wavelengths strongly activates the mouse visual cortex in darkness. (A) Schematic of the recording setup. Red (673 nm), orange (594 nm) or blue (473 nm) laser was delivered via an optical fiber implanted in the visual cortex (VIS, left hemisphere) of awake head-fixed mice. Translaminar Neuropixels recordings were performed in the contralateral VIS in complete darkness. Black dental cement in and around the headplate and blackout adhesive around the optical fiber were used to prevent light emitting through the craniotomy, skull and optical fiber. In addition, eye cups were gently positioned to touch the fur around the eye to further ensure there to be no external activation of the retina. (B) Raster plots of spikes of three individual units in response to ten repetitions of 0.5 s of blue (left), orange (middle) or red (right) laser stimulation and the corresponding average peri-stimulus time histogram (PSTH). Recordings were performed in complete darkness with laser power of 15 or 1mW. Bar plots display the average percentage of the total number of units (n = 761) activated by blue, orange and red laser stimulation across 1, 2.5, 5, 10 and 15mW. Open circles indicate the percentage of responding units for each laser intensity. Error bars represent the sem of the percentage of responders over the five laser intensities. (C) Average spike density functions for all units aligned to the onset (dotted line) of red, orange or blue laser stimulation (n = 761 cells, n = 4 mice) using either 15mW (top) or 1mW (bottom). Grey shaded area indicates sem. (D) Average difference in firing rates (mean ± sem) between a baseline window (0.2 s before stimulation) and a response window (0.5 s after onset of stimulation) for red, orange or blue laser stimulation at different laser power. Asterisks indicate significant differences between red and blue (black) as well as red and orange (gray). (E) Violin plots displaying onset latencies of red, orange and blue laser-evoked responses at 15mW (top) and 1mW (bottom) laser stimulation. Individual data points (circles), median (black circle) and first and third quartiles are displayed (black lines).

Under these conditions we found many individual units that changed their firing rate in response to laser stimulation using 473, 594 and 637 nm at both high (e.g. 15mW) and low (e.g. 1mW) laser intensities (Fig. 1B, Fig.S1, significance assessed by ZETA test, see Methods). Overall, 18-37% of units responded to laser stimulation across all wavelengths and laser intensities tested (Fig. 1B). When averaging across all cells and animals, all wavelengths and stimulus intensities were found to evoke a significant increase in the firing rate of the contralateral visual cortex (e.g. **15mW**: 473 nm baseline = 2.93 ± 0.2 Hz versus laser ON = 5.16 ± 0.33 Hz; 594 nm baseline = 3.41 ± 0.23 Hz versus laser ON = 5.99 ± 0.37 Hz; 637 nm baseline = 3.29 ± 0.2 Hz versus laser ON = 5.99 ± 0.56 Hz; n = 761 units, p < 0.001, **1mW**: 473 nm baseline = 3.28 ± 0.24 Hz versus laser ON = 5.09 ± 0.33 Hz; 594 nm baseline = 3.51 ± 0.23 Hz versus laser ON = 5.04 ± 0.31 Hz; 637 nm baseline = 3.24 ± 0.21 Hz versus laser ON = 6.6 ± 0.44 Hz; n = 761 units, p < 0.001, permutation test, respectively, Fig. 1C and D).

### Red wavelength light causes the strongest modulation of neuronal activit

By pooling across laser intensities we found that red wavelength light reliably evoked responses in the largest fraction of units (range of percentage of responsive units: red, 29-37%; orange, 18-33%; blue, 18-24%; p < 0.05, one-way ANOVA with Tukey-Kramer multiple comparison corrections, Fig. 1B, Fig.S2B). Not only was red wavelength stimulation found to recruit the largest fraction of cells, but also most consistently evoked the largest change in firing rate (Fig. 1D). Already at the lowest laser intensity, 637 nm light illumination led to the strongest change in firing rates when compared to 594 and 473 nm *Δ* firing rate 637 nm = 2.41 ± 0.26 versus *Δ* firing rate 594 nm = 1.01 ± 0.12 and *Δ* firing rate 473 nm = 1.04 ± 0.13, p<0.001, Friedman’s Test with Tukey-Kramer multiple comparison corrections, Fig. 1D).

Consistent with the known scattering properties of these three light wavelengths, we also observed that the onset latencies of the neuronal responses to 637 and 594 nm laser stimulation at 15mW were significantly shorter compared to 473 nm (latency 637 nm = 87.5 ± 4.038 ms and 594 nm = 98.72 ± 4.94 ms versus latency 473 nm = 121.96 ± 6.36 ms, n = 265, 249, 184 units, respectively; p < 0.01, Kruskal-Wallis test with Tukey-Kramer multiple comparison corrections, Fig. 1E). At 1mW laser stimulation intensity, 637 had the shortest latency compared to both 594 and 473 nm (latency 637 nm = 111.4 ± 4.69 ms versus latency 594 nm = 144.3 ± 8.9 ms and latency 473 nm = 141.93 ± 7.66 ms, n = 199, 100, 102 units, respectively; p < 0.01, Kruskal-Wallis test with Tukey-Kramer multiple comparison corrections, Fig. 1E).

### External ambient illumination reduces off-target light-evoked neuronal activation

To determine whether decreasing the sensitivity of the retina could diminish these observed laser-evoked responses in visual cortex we next positioned two monitors in front of the animal and displayed full-field gray iso-illumination at different brightness levels (20, 40 or 80 lux) during laser stimulation (Fig. 2A).

**Figure 2.**
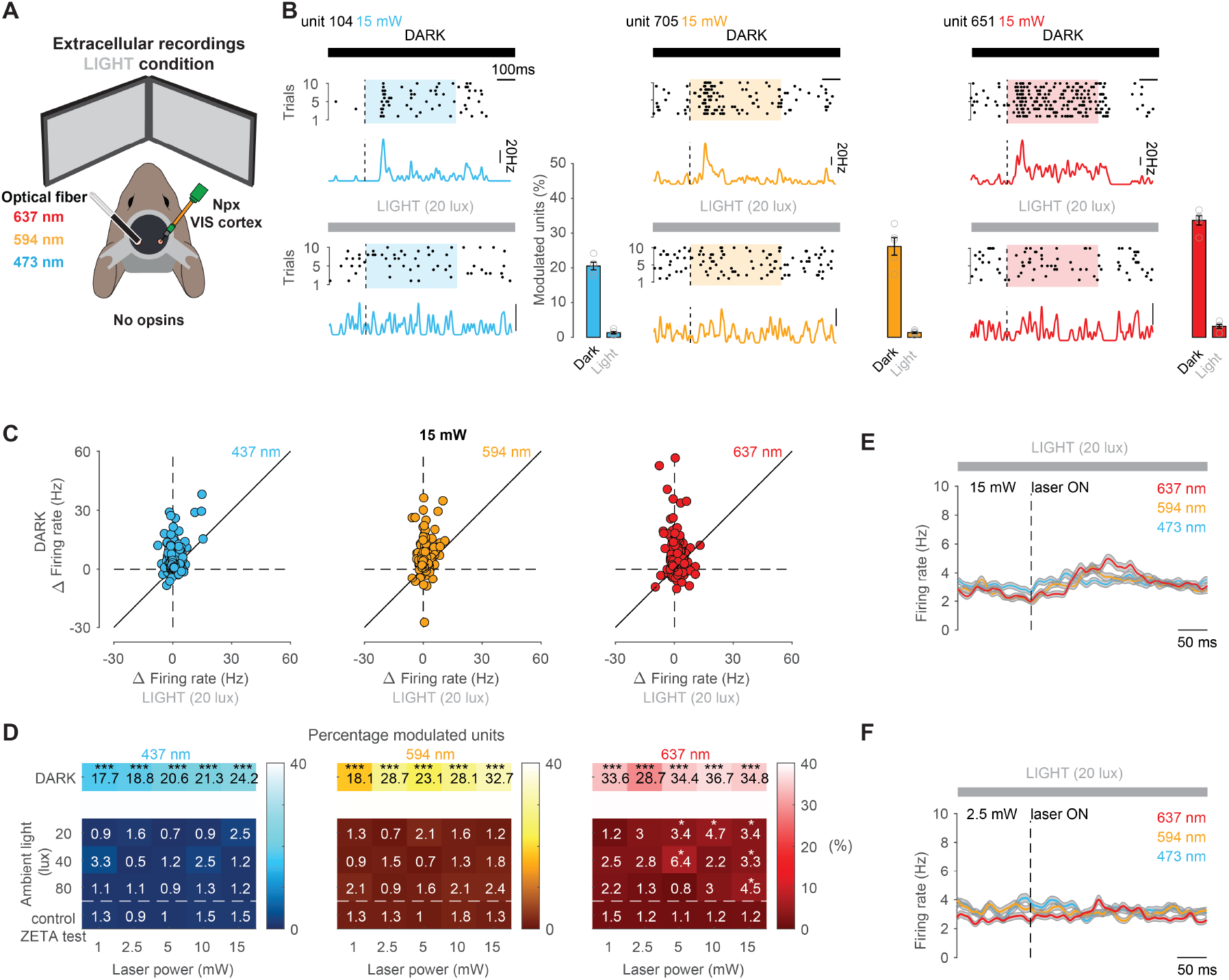
External ambient illumination eliminates internal light-evoked neuronal activation in the visual cortex. (A) Schematic of the recording setup. Red (673 nm), orange (594 nm) or blue (473 nm) laser was delivered via an optical fiber implanted in the visual cortex (VIS, left hemisphere) of awake head-fixed mice. Translaminar Neuropixels recordings were performed on the contralateral VIS. External illumination was delivered via two monitors displaying a full screen iso-illuminant gray image. (B) Raster plots of spikes of three individual units (same cells as in Fig. 1B) in response to ten repetitions of 0.5 s of blue (left), orange (middle) or red (right) laser stimulation and the corresponding average peri-stimulus time histogram (PSTH). Recordings were performed either in complete darkness with laser power of 15 or 1mW or under ambient light conditions (indicated by the grey bar). Bar plots display the average percentage of the total number of units (n = 761 units, n = 4 mice) activated by blue, orange and red laser stimulation across 1, 2.5, 5, 10 and 15mW under darkness or ambient light conditions. Open circles indicate the percentage of responding units for each laser intensity. Error bars represent the sem of the percentage of responders over the five laser intensities. (C) Scatter plot displaying the average laser-evoked firing rate change for those cells that responded under darkness versus ambient light conditions for blue (n = 184), orange (n = 249), red (n = 265) laser stimulations at 15mW. (D) Heatmaps displaying the percentage of units modulated by blue (left), orange (middle) and red (right) laser stimulation under dark (top row) and different ambient light conditions and laser powers. Bottom row displays the false positive percentage for pooled dark and ambient light conditions according to the ZETA test used to assess significant modulation during a control period when laser stimulation was absent. Asterisks indicate significant differences between percentages of modulated units under stimulation periods (rows 1-4) compared to baseline periods without any laser stimulation (row 5). (E) Average spike density functions for all units aligned to the onset (dotted line) of red, orange or blue laser stimulation (n = 761 cells, n = 4 mice) using 20 lux of ambient light brightness and 15mW laser stimulation intensity. Grey shaded area indicates sem. (F) Same as E for 2.5mW laser stimulation intensity.

Under these conditions we found that very few units responded to blue, orange and red laser stimulation at laser intensities between 1-15mW in the presence of 20 lux external illumination (Fig. 2B-D, Fig.S2A and B). At 15mW, even many of the units that were activated by red laser stimulation in darkness significantly reduced their laser-evoked firing in the presence of ambient light (Fig. 2B and C, Fig.S2A, 637 nm *Δ* firing rate dark = 5.36 ± 0.51 versus *Δ* firing rate 20 lux = 0.92 ± 0.16, n = 265 units, p<0.001, two-sided Wilcoxon signed-rank test). At a population level, when comparing baseline firing rates to that recorded during laser stimulation we found no significant increase in neuronal activity at 15mW when using 594 and 473 nm wavelengths in the presence of ambient light (20 lux; 594 nm firing rate baseline = 3.4 ± 0.17 Hz versus firing rate laser ON = 3.9 ± 0.17 Hz; 473 nm firing rate baseline = 3.85 ± 0.19 Hz versus firing rate laser ON = 4.16 ± 0.21 Hz, n = 761; p > 0.05, permutation test, respectively, Fig. 2E). We also found that the proportion of cells that were activated by laser stimulation at 15mW greatly decreased (594 nm ambient darkness: 32.7% or 249/761 versus ambient light: 1.2% or 9/761; 473 nm ambient darkness: 24.2% or 184/761 versus ambient light: 2.5% or 19/761; p < 0.001, Fisher exact test, Fig. 2D). Similar observations were made across all laser intensities and ambient light brightness conditions (Fig. 2D). Generally, the proportion of remaining cells still significantly modulated by 473 and 594 nm laser stimulation under the different laser and ambient light conditions were indistinguishable to the false positive rate detected with the ZETA test during a control period without any laser stimulation (Fig. 2D). While the majority of cells that were activated upon 637 nm laser stimulation at 15mW in complete darkness no longer responded in the presence of ambient light (darkness: 34.8% or 265/761 versus light: 3.4% or 26/761; p < 0.001, Fisher exact test, Fig. 2D), we found that some cells maintained their responsiveness. Indeed, 3.4% of units recorded under ambient light conditions and laser stimulation (at 15mW) was still significantly higher than the false positive proportion detected with the ZETA test during a control period where laser stimulation was absent (light: 3.4% or 26/761 versus control: 1.2% or 9/761; p < 0.001, Fisher exact test, Fig. 2D). This small fraction of cells responded significantly enough to cause an average population increase in activity at 15mW (firing rate baseline = 3.51 ± 0.2 Hz versus firing rate laser ON = 4.22 ± 0.2 Hz, n = 761 units, p < 0.05, permutation test, Fig. 2E). Only at laser powers equal or less than 2.5mW at 20-80 lux ambient light, was the proportion of laser-activated units found to be indistinguishable to the false positive rate (20 lux light: 3% or 23/761 versus control: 1.2% or 9/761; p < 0.001, Fisher exact test, Fig. 2D) and the average population did not significantly increase in activity upon laser stimulation (firing rate baseline = 3.5 ± 0.2 Hz versus firing rate laser ON = 3.92 ± 0.2 Hz, n = 761 units, p > 0.05, permutation test, Fig. 2F, Fig.S2A).

Taken together, these results show that in darkness, optical-fiber stimulation within the mouse brain internally activates the retina and subsequently the visual cortex and that while ambient light can prevent off-target retinal activation over a range of experimentally relevant laser intensities for orange and blue light, it does not completely abolish responses to red light at high laser power.

## 3 Discussion

In this study we found that in darkness, delivery of red, orange and blue light through optical fibers in live brain tissue lacking exogenous opsins is sufficient to robustly activate cortical neurons. Surprisingly, we find that this unintended neuronal activation is triggered even at the lowest levels of laser intensities commonly used in *in vivo* optogenetic experiments (Aravanis *et al*. 2007; Madisen *et al*. 2012; Olsen *et al*. 2012; Pinto *et al*. 2013; Chuong *et al*. 2014; Danskin *et al*. 2015).

In order to exclude light leakage from the craniotomy itself, which potentially could lead to external activation of the eye, we shielded the eyes with eye cups in darkness, covered the dorsal surface of the skull and craniotomy with black dental cement and used a black sleeve around the fiber connector. The simplest explanation for the light-evoked increase in neuronal activity without the expression of exogenous opsins is off-target activation of endogenous opsins. In this scenario, light from the fiber tip propagates through the brain tissue (Yaroslavsky *et al*. 2002; Yizhar *et al*. 2011; Jacques 2013; Danskin *et al*. 2015; Lehtinen, Nokia, and Takala 2022), reaching the back of the eye at intensities that are sufficient to activate retinal photoreceptors and therefore subsequent visual pathways. This idea is supported by periorbital ERG recordings showing that light from implanted optical fibers in the somatosensory cortex can be detected by the retina (Danskin *et al*. 2015). Given the long latencies and the fact that our observed responses are absent when ambient external light is presented to the eye it appears unlikely that these neuronal responses to optical stimulation are due to any direct photoelectric effect on the recording probe (Steinmetz *et al*. 2021).

While it is the case that all three wavelengths tested here led to the unintended activation of neurons, red light caused the most severe effect both in complete darkness and under high laser intensities in ambient light. At first, this seems to be at odds with the wavelength sensitivity of rodent retinal opsins, which peaks at ultraviolet and green spectra and therefore should show little to no activation under red and stronger activation under blue and orange wavelengths. However, it has been recently shown that the rodent retina is more responsive to red light than previously thought, although possibly through the activation of intrinsically photosensitive retinal ganglion cells (Niklaus *et al*. 2020). In addition, rodent opsin spectral properties still show a red-shifted absorption, albeit a 10-100-fold less for 637 nm compared to green (∼500 nm, Bridges 1959; Peirson *et al*. 2018). Therefore, the reduced retinal sensitivity for red could be compensated by less scattering of red light through brain tissue compared to orange/blue light, which yields a sufficient illumination intensity when reaching the retina. Indeed, red-shifted wavelengths scatter less through media including brain tissue, thereby traveling further and preserving higher irradiance compared to orange and blue light (Yaroslavsky *et al*. 2002; Yizhar *et al*. 2011; Jacques 2013; Lehtinen, Nokia, and Takala 2022). In line with this, periorbital ERG recordings have shown that illumination of the retina from an optical fiber implanted in the brain is 50-fold greater when red wavelengths are used compared to shorter wavelengths. Consequently, both the retinal responses measured in a previous study (Danskin *et al*. 2015) as well as the cortical response latencies measured in our study are shorter under red light when compared to orange/blue light stimulation.

In the current study we measured strong neuronal spiking responses to light stimulation within the visual cortex *in vivo* where there was no expression of exogenous opsins. It is however plausible that neuronal activation by brain laser stimulation is more widespread and might affect any brain area connected downstream from the visual system. Furthermore it is expected that off-target retinal activation will have a more profound effect on the subthreshold activity of downstream neurons and this is likely to be even more widespread. It is of utmost importance, therefore, to test optogenetic experimental protocols in the absence of exogenous opsins regardless of the brain area targeted.

Here we show that brain laser stimulation in ambient darkness consistently evokes unintended neuronal activation, while leveraging retinal light-adaptation by exposing the animal to relatively dim ambient light offers a practical and easily implementable solution. We show that this approach is effective under relatively high laser stimulation intensities (particularly for blue/orange wavelengths), while maintaining a low ambient illumination. Factors such as the duration, intensity and wavelength, as well as the fiber implantation site and the proximity of both the stimulation and recording site to the eyes and visual pathway, should be carefully considered prior to establishing optogenetic experiments.

## 4 Methods

### Animals

All experiments were performed on 20-36 weeks old C57BL/6 male mice in accordance with the UK Home Office regulations (Animal (Scientific Procedures) Act 1986), approved by the Animal Welfare and Ethical Review Body (AWERB; Sainsbury Wellcome Centre for Neural Circuits and Behavior) and in compliance with ARRIVE guidelines. Every effort was made to minimize the number of animals and their suffering.

### Surgical procedures

Mice were anesthetized under isoflurane (2%-5%), carprofen or meloxicam was administered (5mg/kg or 1-2 mg/kg, respectively; s.c.) and eyes were protected with eye gel Lubrithal (Dechra). Mice were then fixed to a stereotaxic frame and their body temperature maintained at 37-38 °C. They were implanted with a head plate fixed to the skull using Histoacryl (Braun Medical) and C&B Metabond (Sun Medical). Additionally, we added a layer of black dental cement (kemdent) on top of the Metabond. For the implantation of the optical fiber, a craniotomy of ∼ 1mm radius was drilled over the region of interest using a 0.3mm burr dental drill (Osada Electric). The optical cannula (Newdoon 200 *µm*, 1.5mm, NA 0.37) was then inserted 550 *µm* from the pial surface into the cortex and fixed to the skull using light-cured resin dental cement Relyx Unicem 2 (3M). The skull was then covered with C&B Metabond and subsequently covered with black cement. Animals were allowed to recover for at least 48 hr. On the day of extracellular recording (after habituation to head-fixation that typically took 2-3 sessions of 30 minutes each), animals were anesthetized under isoflurane (2%-5%), their whiskers trimmed (to 2 to 5mm long) to avoid any whisker related proprioception and a small craniotomy (1 × 1mm) was drilled over the brain region of interest using a 0.3mm dental drill. The craniotomy was sealed with silicon kwik-cast (World Precision Instruments) and the animals were allowed to recover for at least 2 hr previous to recording.

### Experimental setup

The visual stimulus was presented on two portable monitors (300 fps; MG300, Magedok). Screens were placed 6 cm from each eye at a 90 degree angle to each other (visual field covered: 75° of elevation and 270° azimuth). In addition, an LED strip (Goove) surrounding the experimental setup was turned on (∼1m away from the animal). The brightness was measured with a Luxmeter (ILM 1337, Isotech) at the position of the eye of the animal. In ambient light experimental conditions, isoluminant gray screens with different brightness (20, 40 and 80 lux) were presented using the Psychophysics Toolbox (Brainard 1997). In complete darkness conditions, the screens and LED strip were turned OFF and two small eye cups were positioned gently on the fur surrounding the eyes. In both light and dark conditions, a light shield was positioned at the base of the optical fiber. A Faraday cage was built around the experimental apparatus to provide both electrical noise isolation and pitch darkness.

### Extracellular recordings

The Neuropixels 2.0 (4 shanks) was first coated with DiI (Molecular Probes, Thermo Fisher Scientific). The probe was then positioned with an angle of 45 degree to the brain surface and the tip inserted 1500 *µm* from the pial surface using micro-manipulators (Luigs and Neumann). The probe was allowed to settle for approximately 30 minutes before recordings. The reference electrode consisted of a Ag/AgCl wire positioned close (<2mm) to the craniotomy. The craniotomy, as well as the reference wire, were covered by Agar 3% prepared in cortex buffer (NaCl 125 mM, KCl 5 mM, Glucose 10 mM, HEPES 10 mM, CaCl2 2 mM, MgSO4 2 mM, pH 7.4). The Agar was then submerged by cortex buffer. The Neuropixels probe was connected to a PXIe card inside a National Instruments chassis. SpikeGLX v20201103-phase30 (https://github.com/billkarsh/SpikeGLX) was used to acquire data. Data was filtered at 300 Hz during or after acquisition.

### Laser stimulation

An optical cord (Newdoon MM200/220 NA 0.37) was used to provide the laser stimulation (637 nm: Thorlabs S4FC637; 594 nm: Coherent OBIS 594nm LS 60mW; 473nm: Coherent OBIS 473nm LX 150mW). In pitch darkness and ambient light with gray screens, 10 laser stimulations were presented in 5 blocks of increasing laser intensities (1, 2.5, 5, 10 and 15mW). Within each block, each laser stimulation lasted 0.5 seconds and was separated by 4 seconds. Each block was separated by 14 s. Laser power was measured after the fiber tip using a digital power meter (PM100D, Thorlabs). Irradiance was calculated using: http://web.stanford.edu/group/dlab/cgi-bin/graph/chart.php.

### Probe localisation

Following silicon probe recordings, animals were deeply anesthetized and transcardially perfused with cold phosphate buffer (PB, 0.1 M) followed by 4% paraformaldehyde (PFA) in PB (0.1 M) and brains left overnight in 4% PFA at 4 °C. Brains were then embedded in 4% agar and imaged using serial two-photon tomography (Ragan *et al*. 2012), using a custom system controlled by ScanImage (Vidrio Technologies) and BakingTray (Campbell 2020). Images were acquired as tiles with 5-10 *µm* axial sampling and 2.27 × 2.27 - 4.13 × 4.13 *µm* pixels, and stitched using StitchIt (Campbell, Blot, and lguerard 2020). Images were then registered to the Allen Mouse Brain Common Coordinate Framework version 3 (CCFv3, 25 *µm* resolution; Wang *et al*. 2020) using brainreg (Niedworok *et al*. 2016; Tyson *et al*. 2022). The atlas data was provided by the BrainGlobe Atlas API (Claudi *et al*. 2020). Probe tracks were confirmed by DiI fluorescence and were traced using brainreg-segment (Tyson *et al*. 2022).

### Spike sorting and analysis

Spike sorting and unit quality control were performed using a fork of the ecephys_spike_sorting pipeline (Siegle *et al*. 2021) modified for SpikeGLX data (https://github.com/jenniferColonell/ecephys_spike_sorting, for Neuropixels 2.0 commit 60c40251ed568fb036b4364e615a261d3afc4800). Raw data acquired with SpikeGLX were filtered with CatGT (https://billkarsh.github.io/SpikeGLX/catgt) and Kilosort2 was run. For single unit analysis only clusters deemed “good” were considered (https://github.com/MouseLand/Kilosort, commit 2a399268d6e1710f482aed5924ba90d52718452a). Double counted spikes were removed (Siegle *et al*. 2021). All units (wide- and narrow-spiking units) were pulled for further analysis. Firing rates during baseline and laser stimulation periods were calculated on a trial-by-trial basis. Spike trains were convolved with a 5 millisecond-wide Gaussian window, to obtain a continuous spike rate.

Laser-responsive units were identified using the ZETA (Zenith of Event-based Time-locked Anomalies) test (Montijn *et al*. 2021). The ZETA test is a parameter-free statistical test that enables testing whether neurons show a time-dependent modulation of their firing rates by an event. The ZETA test was restricted to the period during the 0.5 s laser stimulation and ran with 100 random resamplings to identify which neurons showed significantly light-modulated spiking activity (p < 0.05). To estimate a false positive percentage of the ZETA test, we applied the ZETA test on a control period without any laser stimulation. The ZETA test was restricted to a 0.5 s period and performed ten times before and between each stimulation block.

To calculate the change in firing rate two windows of interest of 200 ms and 500 ms were taken: immediately before (baseline, 200 ms) and immediately after laser stimulation onset (response, 500 ms). The average of the spike rate trace during baseline and response window were then subtracted.

### Statistics

Details of all n and statistical analysis are provided either in the results and/or in the figure legends. Before comparison of data, individual data sets were checked for normality using the Anderson-Darling test in MATLAB 2024. Statistical analyses were performed using MATLAB 2024. Asterisks indicate significance values as follows: *p<0.05, **p<0.01, ***p<0.001

## 5 Author Contributions

S.W., M.F.V. and T.W.M conceived the project. S.W. and M.F.V. performed all experiments. S.W. analyzed all data. S.W., M.F.V. and T.W.M interpreted and discussed all data and wrote the manuscript.

## Acknowledgments

The authors are grateful to the support staff of the Neurobiological Research Facility at Sainsbury Wellcome Centre and Manuel Teichert for comments on the manuscript. T.W.M., M.F.V. and S.W. are funded by The Wellcome Trust (214333/Z/18/Z; 090843/F/09/Z).

## Data, Materials, and Software Availability

Analysis code and structure of processed data will be deposited upon acceptance of this manuscript. Data available on request.

## Notes

### Competing Interest Statement

The authors have declared no competing interest.

### Summary of Updates

- There was a mistake in the abstract as it stated "yellow" and not "orange" light - in the Introduction we changed "rodent" to "rodents"

