## Supporting information for "Overcoming off-target optical stimulation-evoked cortical activity in the mouse brain *in vivo*"

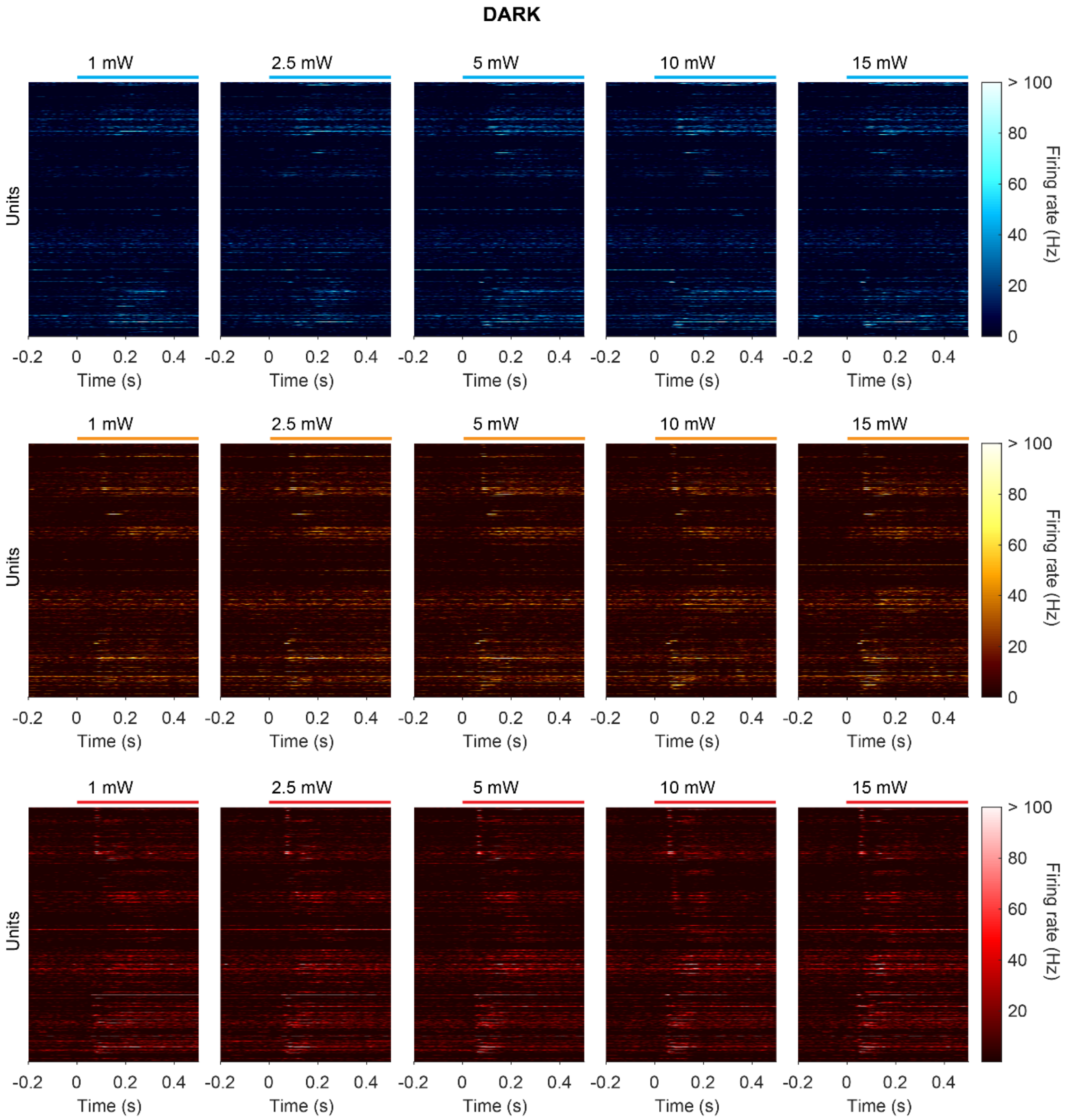


**Fig.S1: Laser stimulation in the absence of exogenous opsins during extracellular recording in the contralateral visual cortex in darkness.** Spike density functions for all units aligned to the onset (stimulation window displayed on top) of blue (top), orange (middle) or red laser stimulation (n=761 cells, n=4 mice) for five laser stimulation intensities in complete darkness

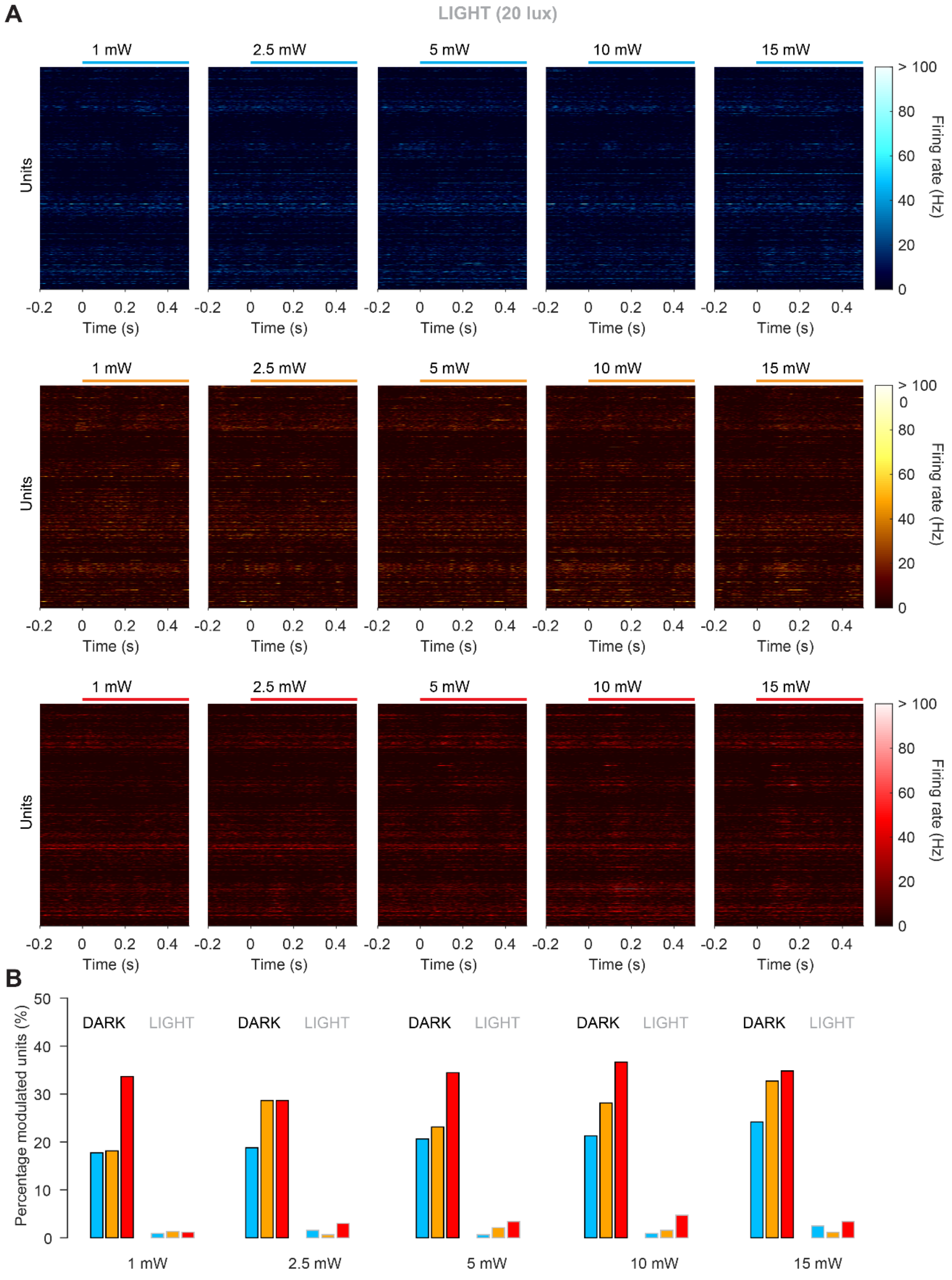


**Fig.S2: Laser stimulation in the absence of exogenous opsins during extracellular recording in the contralateral visual cortex under ambient light conditions.** (A) Spike density functions for all units aligned to the onset (stimulation window displayed on top) of blue (top), orange (middle) or red laser stimulation (n=761 cells, n=4 mice) for five laser stimulation intensities under ambient light of 20 lux. (B) Percentage of modulated units for blue, orange and red laser stimulations at different laser powers in darkness or under 20 lux of ambient light.
